# Great tits show serial reversal learning in the perseverance phase but not in the new learning phase

**DOI:** 10.1101/2025.03.27.645707

**Authors:** Ernő Vincze, Anders Brodin

## Abstract

An animal possessing reversal learning ability is capable of unlearning a previously learned association between a stimulus and a reward and learning a new contingency. It is a form of behavioural flexibility which can be advantageous in changing environments. Serial reversal learning occurs when an animal’s performance improves over repeated reversals of contingencies. In this study, we tested the serial reversal learning ability of great tits (Parus major) in an aviary experiment where they could choose between two laterally positioned locations marked with different symbols. One of the two locations contained hidden food reward, and the side that was rewarding and unrewarding was reversed several times for each bird. We divided the learning process after each reversal into two phases, the perseverance phase and the new learning phase, quantified by the number of visits before and after the first visit to the newly rewarding location respectively. We found that the length of the perseverance phase significantly decreased over repeated reversals. However, there was no corresponding decrease in the length of the new learning phase. This suggests that perseverance and new learning are separate cognitive processes, and that the former may be less challenging than the latter. Initially the birds also showed a colour preference for yellow over blue, but this did not affect their reversal learning ability. The high behavioural flexibility of great tits may help explain their success in exploiting challenging environments.

**Highlights:**

- Perseverance of great tits decreases in a serial learning task
- New learning does not become faster after multiple reversals
- Preference for yellow colour over blue in the initial choice

## Introduction

Cognition, i.e. the ability to acquire, process, memorize and use information from the environment (Shettleworth, 2010) can help animals cope with environmental challenges. Cognition is related to behavioural plasticity, because high cognitive ability allows animals to modify their behaviour based on the environmental information they gain. One of the most studied cognitive abilities in animals is associative learning, i.e. the ability to link a neutral stimulus with a biologically relevant stimulus, such as a food reward. This is the very basis of classical conditioning experiments that have been performed for over a century (Pearce & Bouton, 2001). Though most studies on learning abilities have been performed in captivity, more recent studies have also investigated it in wild populations (Morand-Ferron, Hamblin, Cole, Aplin, & Quinn, 2015). Although simple associative learning is very useful in a relatively stable environment where certain stimuli are strongly connected with certain rewards, memorizing a permanent association will be disadvantageous in a changing environment where the connection between various stimuli and rewards may be temporary. For example, food sources may be depleted or only available at certain times of the year. In such cases, animals will benefit from the ability to replace old associations with new, relevant ones.

This kind of behavioural flexibility is usually studied in a type of experiments known as reversal learning tests, first developed by Williams (1942). These tests usually start with an associative learning stage during which animals need to make a choice between two options, one rewarding and one unrewarding. After an animal has learned which option is rewarding and which is unrewarding, the conditions are reversed so that the previously unrewarding option becomes rewarding, and vice-versa. Hence the animal must learn this new contingency. Typically, immediately after the reversal, the animal keeps choosing the previously rewarding option before checking the previously unrewarding one. This behaviour is usually referred to as “perseveration” or “perseverance” (Biondi, Medina, Bonetti, Paterlini, & Bó, 2024; Hervig et al., 2020). Following this perseverance phase, the animal begins to learn the new contingency, associating the previously unrewarding option with the reward. Incorrect choices during this new learning phase are sometimes referred to as “regressive errors” (Biondi et al., 2024). The neurochemical mechanisms during the perseverance phase and the new learning phase are different (Hervig et al., 2020), suggesting that “unlearning” the old contingency and learning the new contingency are distinct cognitive processes. The relationship between associative learning and reversal learning is unclear. Some studies suggest a negative correlation between associative and reversal learning, likely because a stronger associative learning should result in greater perseverance and therefore slower reversal learning, a mechanism known as proactive interference (Hermer, Murphy, Chaine, & Morand-Ferron, 2021). Other studies, however, found a positive relationship between associative learning and reversal learning, suggesting that the two are part of a general learning ability (van den Heuvel, Quinn, Kotrschal, & van Oers, 2023).

A handful of studies go one step further, and instead of just doing one reversal, they perform multiple reversals (Cauchoix, Hermer, Chaine, & Morand-Ferron, 2017; Hermer, Cauchoix, Chaine, & Morand-Ferron, 2018). These serial reversal learning studies are a good way to test whether animals perform consistently in these tasks (repeatability) and/or improve over repeated reversals. If reversal learning performance improves over these repeats, this suggests that the animals understand the concept of reversals. This should help them to make better decisions in a changing environment (Hermer et al., 2018; Izquierdo, Brigman, Radke, Rudebeck, & Holmes, 2017). Improvement during serial reversal learning has been demonstrated in several taxa, both in specialized nectar-feeding species like bats (Chidambaram, Wintergerst, Kacelnik, Nachev, & Winter, 2024) and bumblebees (Strang & Sherry, 2014) where the flowers that have been emptied by foragers get replenished over time, and in generalist foragers like corvids (Bond, Kamil, & Balda, 2007) and great tits *Parus major* (Cauchoix et al., 2017; Hermer et al., 2018). For generalists, reversal learning can be particularly useful when exploiting anthropogenic resources, for example visiting bird feeders that may be filled up irregularly. There is evidence that the change in perseverance and the change in learning speed over repeated reversals are driven by different neurochemical mechanisms (Hervig et al., 2020). Despite this, the above studies (except for Chidambaram et al. 2024) did not distinguish between the perseveration phase and the learning phase.

In this study we aimed to test the performance of great tits in a serial reversal learning task, i.e. whether their perseverance and/or their learning becomes shorter with each reversal. This species performs well in various cognitive tasks (Johnsson & Brodin, 2019; Preiszner et al., 2017), including associative learning (Morand-Ferron et al., 2015), single reversal learning (Amy, van Oers, & Naguib, 2012; van den Heuvel et al., 2023) and serial reversal learning (Cauchoix et al., 2017; Hermer et al., 2018), but it has not been tested yet whether the improvement in their performance is a consequence of shorter perseverance or faster learning. In addition to this question, we also tested whether the birds show any preference for lateral side, colour or geometric shape when initially having to choose between two stimuli, and whether laterality, colour, shape of the symbol, or properties of the birds such as sex and age affect their perseverance and learning speed.

## Methods

### Subjects and housing

We captured great tits from 3 locations within the city of Lund, Sweden (Site 1: 55.7144 N, 13.2069 E; Site 2: 55.7226 N, 13.1947 E; Site 3: 55.6976 N, 13.2472 E) between October 2022 and February 2023. All three capture sites were in urban habitats, because earlier studies suggest that urban animals habituate more easily to captive conditions (Vincze et al., 2016) and that in this species urban birds perform better in cognitive tasks than their rural conspecifics (Preiszner et al. 2017; Grunst et al. 2019, but see Vincze et al. 2024). We ringed the birds with one numbered metal ring and one or two plastic colour rings for individual identification in the lab. We estimated the age and sex of the birds using plumage characteristics. In our sample of 17 birds, there were 3 juvenile and 6 adult females, and 1 juvenile and 7 adult males. An 18^th^ bird, an adult male from the second batch, was captured but released as it did not pass the training phase (see below) in 5 consecutive days.

The birds were housed separately in 55 × 56 × 36 cm cages that we had positioned on shelves in an enclosed compartment along a wall in an aviary room, adjacent to our experimental room. This room had lighting with an outdoor light spectrum and computer-controlled light and temperature regimes. In mornings and evenings, an automatic one-hour dimming function simulated dawn and dusk, roughly following outdoor day length patterns. We kept the temperature constant at 14°C, which is a temperature that works well in this type of experiment (Brodin & Urhan, 2014, 2015; Isaksson, Urhan, & Brodin, 2018; Vincze et al., 2024). From their home cages, each bird had close visual contact with at least one other bird. The birds had *ad libitum* access to a diet of mixed seeds (sunflower, millet, peanuts), but no animal food, because great tits tend to be more successful in cognitive tasks when kept on a seed diet (Davidson et al., 2020), presumably due to higher motivation level. The birds were also provided water that was changed daily and enriched with a commercial vitamin supplement for birds. We cleaned the cages every day. Before the experiment started, the birds were allowed to acclimatise to the cage and aviary for two days. According to our earlier experiences, this is sufficient for the birds to acclimatise to captivity (Brodin & Urhan, 2014, 2015; Isaksson et al., 2018; Vincze et al., 2024).

### Ethical note

Before each testing session, we visually inspected the birds and made sure that they were in good condition. We avoided handling the birds during the experimental sessions to minimise stress. Instead, we always moved the birds between the aviary and the test room together with their home cage. After the finalization of the experiment, the birds were visually inspected again, and then released at the same site as where they had been captured. The entire experimental protocol was performed in accordance with the permit 4716/2018 (2021-2022) from the Malmö-Lund ethical permit board for animal experiments.

### Experimental setup

The experimental cage (Figure S1) was set up in a test room adjacent to the aviary room. It was positioned on a desk facing an observation booth with a one-way window where the observer could not be seen by the birds. The birds were always taken to the test room together with their home cage to reduce handling stress. The entrance of the home cage could be connected to the main compartment of the experimental cage with a short tunnel. From this main compartment, the bird could access two smaller compartments that were separated from each other. In each of the two compartments, a wooden box (9 cm tall, 7.5 cm wide, 6 cm deep) was positioned. The boxes had a symbol on top: either a full circle (O) or a four-point star (X), coloured either yellow or blue. Each symbol was always paired with the opposite colour and shape: a yellow O with a blue X, and a blue O with a yellow X. The position of the symbols was kept constant for each bird, but differed between individual birds (Table S1).

The birds all participated in a training period during which they familiarized with the experimental cage. The training started three days after capture, and consisted of two phases. In the first phase, we placed five mealworms in a small plastic dish near each of the two boxes where they were visible for the birds (on the top of the boxes for the first five birds; next to the boxes on the floor for the others to reduce the risk of the birds pushing down the dish and spilling the worms), and allowed the birds for 1 hour per day to explore the cage and find the mealworms. This phase was completed when the bird ate some of the mealworms near both of the boxes, which took 1 to 6 days (mean ± SD = 2.12 ± 1.36). On the first day of this stage we did not remove the food from the birds’ home cage, but on the second day and all consecutive days we removed it directly before the start of the training session. In the second phase, the dishes with the mealworms were placed behind the boxes where they were not visible for the birds; the birds were once again allowed to explore the cage and feed for 60 minutes. For the first five birds, there was a third training phase, where the food was placed under the box like in the test sessions. Both of these phases were only 1 day long as the bird always ate the worms, which suggests they learned quickly where to search for the worms. Therefore, the third training phase was deemed unnecessary and removed from the protocol for the other 12 birds to make the training more time-efficient.

Following the training, each bird participated in one test session per day, which started sometime between 9:58 and 15:22. To increase the motivation to feed, all food had been removed from the tested bird’s cage 30 minutes before the test sessions. Before bringing the bird into the test room, the observer first placed one mealworm in a plastic dish on one side, and an empty plastic dish to the other. The observer would then turn off the lights in the test room, move the bird’s home cage to the test cage and attach the tunnel to the cage’s door. Following this, the observer turned on the lights, started the video recording, opened the tunnel, and then entered the observation booth, becoming hidden from the bird. When the bird entered one of the compartments in the cage’s front, the observer noted which side the bird went.

If the bird chose the rewarding side (“correct choice”), then, after the bird left the compartment and ate the worm but before it could return to either compartment, the observer would leave the observation booth, which typically resulted in the bird getting flushed back to its home cage. The observer would then take a mealworm in one hand out of the bird’s sight, then walk to the cage’s front clenching both hands so that the bird would not know which hand the mealworm was in. The observer would then put both hands simultaneously in the compartments, touching both plastic dishes under the boxes at the same time, and put the mealworm in one dish while leaving the other empty. This procedure ensured that the bird could not directly see, only infer, in which compartment the mealworm had been placed. If the bird chose the unrewarding side (“incorrect choice”), then the observer would leave the observation booth immediately flushing the bird and hence ensuring that it couldn’t enter the compartment on the other side. Following this, he would walk to the test cage with clenched hands, put his hands in the compartments and handle the plastic dishes in the same way as above, but would not place any mealworm in either of the dishes (as there was already one mealworm on the rewarding side). In both cases, the observer would then return to the observation booth, and wait until the bird entered one of the compartments, and note which side the bird chose. This procedure was repeated until one of the following conditions applied: a) the bird went to the rewarding side and took the mealworm 4 times; b) the bird went to the unrewarding side 6 times; c) the bird went to the rewarding side but did not take the mealworm; or d) the bird did not go to either side for 10 minutes. We henceforth refer to each time a bird entered one of the compartments as “choices” or “attempts”.

The worm was hidden on the same side until the bird chose the correct side in 9 out of 10 attempts, which is the same learning criterion that earlier studies (Cauchoix et al., 2017; Hermer et al., 2018) used for this species. These 10 attempts would span over 3 test sessions, as each session was terminated after 4 correct choices in order to avoid satiation. When the learning criterion was met, the roles of the sides were reversed: the mealworm was placed in the dish on the previously unrewarding side, whereas the dish was left empty in the previously rewarding side. Note that with the reversal the boxes’ position remained the same, i.e. the same symbol remained on the left and the right side, and it was only the position of the reward that was reversed. The reversal was made immediately after the bird had met the learning criterion (i.e. made the 9^th^ correct choice out of 10) if it was the 1^st^, 2^nd^ or 3^rd^ correct choice on a given day. If the learning criterion was met during the 4^th^ choice on that day, the test session was terminated as usual. The next day would then start with unreversed conditions (i.e. the worm being on the same side as the previous day), and the reversal was made after the 1^st^ correct choice of that day. It was done this way to ensure that the bird always had a short-term experience with the worm being on one side before the reversal was made. We continued this test for 13 to 26 (mean: 20.59) days for each bird, which, depending on their learning speed, resulted in 59 to 142 (mean: 116.47) attempts and 3 to 6 (mean: 4.65) reversals. The experiment was terminated after the first correct choice following the last reversal. All test sessions were recorded on video (type: Toshiba Camileo S20), serving as a backup for the data from the live observation.

### Statistical analyses

First, we tested with chi-squared tests for given probabilities whether the birds at the start of the test showed any kind of bias toward the stimuli at the box with or without the worm, the left or right side, the X or O shape, and the yellow or blue colour, i.e. whether the probability of going to one box or the other differed from 50%. We tested these four explanatory variables in separate chi-squared tests due to the sample size (N=17) not being enough to include all four of them in the same statistical model.

We then tested two aspects of reversal learning: (i) **perseverance**, which was defined as the number of attempts until the first correct choice (i.e. how many times did the bird go to the unrewarding side before going to the rewarding side for the first time); (ii) **new learning**, the number of attempts following the first correct choice until the bird met the learning criterion (went 9 out of 10 times to the rewarding side). We included only the attempts following the first reversal in this variable, because the attempts before the first reversal show simple associative learning rather than reversal learning. However, on Figure S2 we show all attempts, both before and after the first reversal. The two response variables did not correlate with each other (Pearson’s correlation: R = 0.006, t = 0.043, df = 58, P = 0.966), indicating that they are statistically independent. Both variables were tested in generalized linear mixed-effects models with negative binomial distribution, using the „glmmPQL” function of the R package MASS (Ripley et al., 2013). In both models, we included the following covariates: the number of reversals until the trial (to test whether perseverance and/or learning is getting shorter with each trial, which would be an indicator of serial reversal learning) and the total number of reversals for each bird (to control for the fact that birds who were faster learners got more reversals overall). We also included the sex and age of the bird as well as side, colour, and shape where the worm was hidden in a given trial as fixed factors. Additionally, we included site and bird ID as random factors to control for a possible among-site and among-individual variation and test repeatability (see below). We considered variables with P < 0.05 as statistically significant.

We also tested whether the length of the perseverance phase and the learning phase was repeatable within site and within individual. We squared the SD estimates for our random factors (site, individual ID nested within site) and the residual SD to get the three variance components. Then, based on the recommendation of Araya-Ajoy et al. (2015), we estimated within-site repeatability as dividing the variance component for site with the sum of the three variance components, and within-individual repeatability by dividing the sum of the variance components for site and individual ID with the sum of the three variance components. The reason for this approach is that among-individual variation includes both differences between individuals within a site and differences between individuals from different sites.

The full R code for the statistical tests is provided in the supplementary material (S1) along with the code used for creating the figures. The R package “gplots” (Warnes et al., 2016) was used for creating Figure 1.

**Figure 1.**
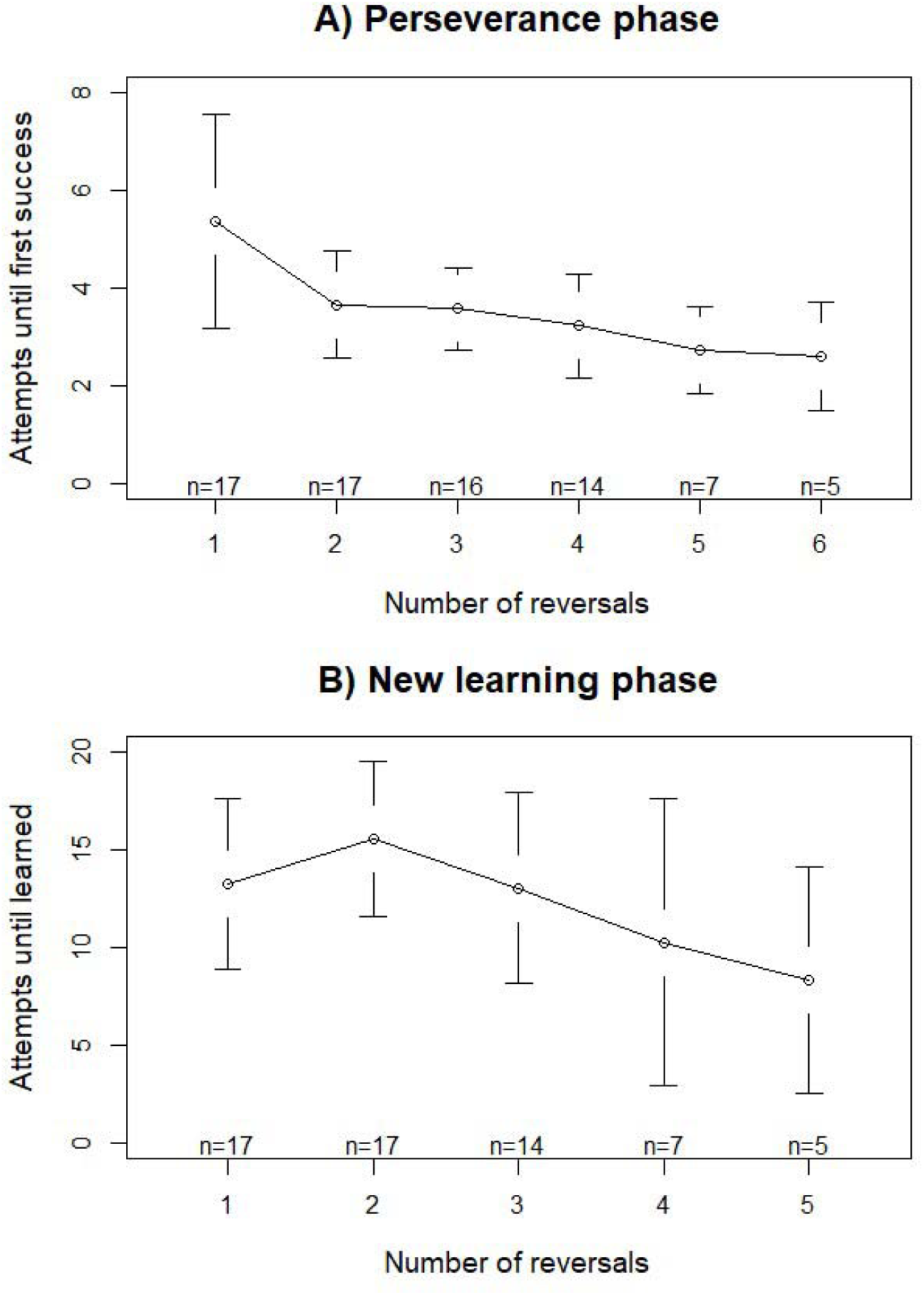
Number of attempts (means + confidence intervals) (A) from the reversal until the first successful attempt (perserverance phase) and (B) from the first successful attempt until meeting the learning criterion (new learning phase), following each reversal. The number of individual birds is indicated at the bottom. See also Figure S2 for the individual reaction norms.

## Results

### Initial choice

At the start of the test, 10 out of 17 birds chose the rewarding side on their first attempt (χ^2^_df=1_ = 0.529, P = 0.467). All the 7 birds that started with choosing the unrewarding side chose the rewarding side on their second attempt. On their first attempt, 9 birds chose the left side and 8 the right side (χ^2^_df=1_ = 0.059, P = 0.808); 12 birds chose the O and 5 the X shape (χ^2^_df=1_ = 2.882, P = 0.090); and 14 birds chose the yellow and 3 the blue colour (χ^2^ = 7.118, P = 0.008), meaning that 9 out of 9 bird chose the yellow O over the blue X and 5 out of 8 birds chose the yellow X over the blue O.

### Perseverance

The number of preceding reversals had a significant effect on the number of attempts until the first correct choice, with fewer attempts after each reversal (Table 1A, Figure 1A, Figure S2). The total number of reversals, sex, age, side, colour and shape had no significant effect (Table 1A). The behaviour was not repeatable within site or within individual (Table 1A).

**Table 1.** Summary of the generalized linear mixed-effects models. Statistically significant fixed effects are highlighted.

| **A) Perseverance (Attempts between reversal and first success)** |  |  |  |  |  |
| --- | --- | --- | --- | --- | --- |
| **Fixed effects** | **Estimate** | **± SE** | **DF** | **t-value** | **P-value** |
| Intercept | 1.517 | ± 0.343 | 55 | 4.426 | 0.000 |
| **Preceding reversals** | **-0.153** | **± 0.048** | **55** | **-3.203** | **0.002** |
| Total reversals | 0.050 | ± 0.067 | 11 | 0.752 | 0.468 |
| Sex (male vs female) | -0.048 | ± 0.134 | 11 | -0.357 | 0.728 |
| Age (young vs adult) | -0.027 | ± 0.163 | 11 | -0.168 | 0.870 |
| Side (right vs left) | 0.174 | ± 0.133 | 55 | 1.310 | 0.196 |
| Colour (yellow vs blue) | -0.156 | ± 0.133 | 55 | -1.174 | 0.245 |
| Shape (X vs O) | 0.003 | ± 0.133 | 55 | 0.026 | 0.979 |
| **Random effects** | **SD** | **V** | **R** |  |  |
| Site | <0.001 | <0.001 | <0.001 |  |  |
| Bird ID in Site | <0.001 | <0.001 | <0.001 |  |  |
| Residual | 0.482 | 0.233 |  |  |  |
| **B) New learning (attempts between first success and learning criterion)** |  |  |  |  |  |
| **Fixed effects** | **Estimate** | **± SE** | **DF** | **t-value** | **P-value** |
| Intercept | 3.894 | ± 0.453 | 39 | 8.591 | 0.000 |
| Preceding reversals | 0.000 | ± 0.069 | 39 | 0.006 | 0.995 |
| **Total reversals** | **-0.311** | **± 0.086** | **11** | **-3.606** | **0.004** |
| Sex (male vs female) | 0.179 | ± 0.158 | 11 | 1.133 | 0.281 |
| Age (young vs adult) | -0.043 | ± 0.207 | 11 | -0.206 | 0.841 |
| Side (right vs left) | -0.090 | ± 0.154 | 39 | -0.586 | 0.561 |
| Colour (yellow vs blue) | 0.156 | ± 0.155 | 39 | 1.003 | 0.322 |
| Shape (X vs O) | 0.091 | ± 0.155 | 39 | 0.587 | 0.561 |
| **Random effects** | **SD** | **V** | **R** |  |  |
| Site | 0.213 | 0.045 | 0.138 |  |  |
| Bird ID in Site | <0.001 | <0.001 | 0.138 |  |  |
| Residual | 0.532 | 0.283 |  |  |  |

### New learning

Out of the explanatory variables, only the total number of reversals had a statistically significant effect on the number of attempts until the bird met the learning criterion (Table 1B), whereas number of preceding reversals (Figure 1B, Figure S2), sex, age, side, colour or shape) had no statistically significant effect (Table 1B). The behaviour was repeatable within site (suggesting that there were differences between sites), but within-individual repeatability was not higher than within-site repeatability, suggesting that the behaviour of birds from the same site did not differ from one another.

## Discussion

We found that with each reversal, the great tits’ perseverance decreased, i.e. on average they approached the previously rewarding box fewer times before checking the previously unrewarding box, although there was considerable variation between individuals in the learning patterns (Figure S2). This suggests that the birds got better at understanding that the previously unrewarding box can become rewarding again. There have been earlier studies (Cauchoix et al., 2017; Hermer et al., 2018) that also supports the idea that great tits are capable of serial reversal learning, which suggests good cognitive flexibility. This flexibility, combined with the good performance in other cognitive tasks (Johnsson & Brodin, 2019; Sasvári, 1979), may explain why the great tit is particularly successful in exploiting anthropogenic habitats.

Interestingly, despite the serial learning effect in the perseverance phase, there was no significant decrease over sessions in the new learning phase, i.e. how many attempts it took until they stopped going to the unrewarding side. This is in line with an earlier study on nectar-feeding bats, where they also reported that the strongest change during serial reversal learning was a decrease in perseverative errors (Chidambaram et al., 2024). There are several possible explanations why we found change in perseverance but not in learning. First, it is likely that meeting the criteria for completing the perseverance phase was an easier task than meeting the criteria for completing the new learning phase. Understanding that “if there is no reward on one side, check the other side” is relatively simple compared to “if there is reward on one side, stop checking the other side”. Reversal learning studies on other species also suggest that acquiring a new association is more challenging than inhibiting a previously learned one, therefore the animals make more errors in the new learning phase than in the perseverance phase (Biondi et al., 2024).

Second, some studies suggest that between the perseverance phase and the new learning phase there is a “random search” phase, during which the animal has overcome the initial perseverative errors but has not started acquiring the new association yet, and therefore has no expectation regarding the reward’s location and chooses randomly between the two options (Hervig et al., 2020). The point between the random search phase and the actual new learning phase is very hard to determine; the usual practice is to look at the proportion of correct and incorrect choices in a “rolling window” (e.g. the last 10, or 30 choices) and determine a threshold (e.g. 7 out of 10 or 20 out of 30 correct choices; the latter was used by Hervig et al. 2020). When the animal crosses this threshold, it is assumed that the actual learning has begun. However, as this threshold is rather arbitrary, we opted to treat the random search phase as part of the learning process, and distinguished only between the perseveration phase and the learning phase, similarly to Biondi et al. (2024). Nevertheless, it is possible that the lack of improvement in the new learning phase is due to an increase in the random searching phase following the second reversal, when the initial rewarding side became rewarding again. Instead of understanding that a second reversal has happened, the birds may have interpreted the new contingency in a way that that the reward can be under either of the boxes, and therefore searched randomly for a longer time. This idea is supported by the fact that the learning phase, on average, was the longest after the second reversal (Figure 2, Figure S3).

Finally, due to the requirement that the observer needed to be present and actively hiding the reward before each attempt, we could only test each bird for a limited amount of time, resulting in a relatively low number of reversals for each bird (2 to 6, on average 4.65). By comparison, Hermer et al. (2018) and Cauchoix et al. (2017) found serial reversal learning in great tits by collecting data using an automatized operant box, allowing for up to 18 reversals per bird in the former and 48 reversals per bird in the latter. It is possible that if our design had allowed for a higher number of reversals per bird, then we would have also detected an improvement in the new learning phase.

In addition, we also found that in the new learning phase there was some variation between the different capture sites, as indicated by the low but non-negligible within-site repeatability. Although we only included site as a random effect to avoid over-parametrizing our models, if we look at the number of attempts in the new learning phase, it is higher in one of the sites (Site 3: mean ± SD = 17.17 ± 7.26) than in the other two (Site 1: mean ± SD = 10.80 ± 7.43; Site 2: mean ± SD = 12.29 ± 8.36). A handful of studies suggest that habitat urbanization affects reversal learning abilities, though the results are inconclusive: some found animals more urbanized areas (Biondi et al., 2024), or areas with more intensive anthropogenic change (Vardi & Berger-Tal, 2022) are better at reversal learning, whereas other studies suggest better reversal learning in rural individuals (Federspiel et al., 2017) or no difference between urban and rural conspecifics (Audet, Ducatez, & Lefebvre, 2016). However, in our study, all three sites were green areas in an urban environment, therefore it is unlikely that this variation was caused by habitat urbanization. Furthermore, this slowest-learning group was also the last of our four batches (see Figure S2), captured in February, therefore it is possible that what we detected here was a seasonal effect rather than a habitat effect.

A limitation of our study was that each bird, regardless of learning speed, was in captivity for a similar amount of time. As a consequence, the birds that were faster learners would get more reversals compared to the slower learners. This, in theory, could lead to among-individual variation leading to a change in mean behaviour that resembles serial reversal learning, if the mean number of attempts after the first few reversals is calculated from both fast- and slow-learning birds but following the later reversals it is calculated only from fast learners. However, we believe that this is not the case in our results, for three reasons. First, we found the serial reversal learning pattern only in the relatively short perseverance phase, but not in the relatively long new learning phase that affected the amount of time between two reversals to a greater extent. Second, we controlled for the total number of reversals for each bird by including it in our models as an explanatory variable, which had a statistically significant effect in the new learning phase, but not in the perseverance phase where the significant serial reversal learning effect was found. Finally, if there was significant variation in the learning speed of the birds (i.e. there were faster and slower individuals), then we would have found high within-individual repeatability. Instead, we found that individual identity explained virtually none of the total variance in the perseverance phase and only the across-site (and not the within-site) variation in the new learning phase. Taken together, these all suggest that the decrease in perseverative errors that we found is not due to among-individual variation in the number of reversals.

Additionally, we also found that at the beginning of the test, before the reversal, the majority of great tits (82%) chose the box that was marked with a yellow symbol, especially when the yellow circle was paired with the blue star. This may partly be because yellow was brighter and had a stronger contrast with the black background, making it more interesting or attractive for the birds to investigate compared to the blue symbol which blended more into the black background. However, great tits may also be interested in the colour yellow because their plumage is partly yellow, and this carotenoid-based colouration is suggested to be an indicator of their nutritional condition (Senar, Figuerola, & Domènech, 2003), whereas he colour blue does not appear in the plumage of great tits and therefore it may be less attractive to them. It may be interesting to test this idea with the closely related blue tit (*Cyanistes caeruleus*), a species where blue and yellow colours are both present in plumage, which are often assumed to have signalling functions (but see Parker 2013). Note that this bias for colour has not affected reversal learning, as colour had no significant effect either on perseverance or on learning.

Overall, we conclude that great tits are capable of serial reversal learning, but it manifests mainly in the perseverance phase rather than the new learning phase. When analysing reversal learning, it is important to treat perseverance and new learning as two separate cognitive processes. For the future, it would be interesting to investigate the potential habitat effects and between-species differences, and also separate the effect of different cues (location, colour, shape) that the birds may use to identify the reward’s location.

## Supporting information

Table S1, captions for Figures S1 and S2, and R code

## Data availability

Our data and the R code for statistical analysis and figure creation is available in the Supplementary Material.

