## Supplementary material for "Great tits show serial reversal learning in the perseverance phase but not in the new learning phase": Table S1, captions for Figures S1 and S2, and R code

**Table S1.** The list of all birds participating in the experiment, with capture site, sex, age, the starting date of the experiment, the shape and colour of the symbol on the feeder on the left and the right side, and the starting side (where the food was initially hidden before the first reversal).

|  |  |  |  | **Left** | | **Right** | |  |
| --- | --- | --- | --- | --- | --- | --- | --- | --- |
| **Site** | **Sex** | **Age** | **Start date** | **Color** | **Shape** | **Color** | **Shape** | **Start** |
| 1 | male | adult | 2022.10.12 | yellow | o | blue | x | left |
| 1 | female | adult | 2022.10.11 | yellow | x | blue | o | left |
| 1 | male | adult | 2022.10.10 | yellow | x | blue | o | right |
| 1 | female | adult | 2022.10.11 | blue | o | yellow | x | right |
| 1 | male | adult | 2022.10.11 | blue | x | yellow | o | right |
| 2 | female | young | 2022.11.17 | yellow | o | blue | x | left |
| 2 | male | adult | 2022.11.23 | blue | o | yellow | x | left |
| 2 | female | adult | 2022.11.17 | blue | x | yellow | o | left |
| 2 | male | young | 2022.11.16 | yellow | o | blue | x | right |
| 1 | female | young | 2023.01.16 | blue | o | yellow | x | left |
| 1 | female | young | 2023.01.17 | blue | x | yellow | o | right |
| 1 | female | adult | 2023.01.15 | yellow | x | blue | o | right |
| 3 | male | adult | 2023.02.13 | blue | o | yellow | x | right |
| 3 | female | adult | 2023.02.13 | yellow | o | blue | x | right |
| 3 | male | adult | 2023.02.15 | blue | x | yellow | o | left |
| 3 | female | adult | 2023.02.14 | blue | x | yellow | o | right |
| 3 | male | adult | 2023.02.13 | yellow | x | blue | o | left |

**Figure captions:**

**Figure S1:** Schematic drawing (A) and photos (B, C) of the experimental cage. The cage’s base was 100×100 and it is 63 cm high. The front of the cage, facing the observer, was a transparent plastic window, with a wooden door on the bottom. The two lateral sides and the back of the cage were covered with wire mesh. The back of the cage was also wire mesh, but also covered with a layer of cardboard to make it opaque. In the centre middle of the backside of the cage, there was a slide door with a short tunnel, which could be connected to the entrance of the bird’s home cage. The experimental cage was divided into three compartment: a main compartment in the back (with a long perch across) and two smaller compartments in the front, separated with a cardboard panel from each other and with plastic mesh from the main compartment with plastic mesh. On the plastic mesh, there were two holes as far from each other as possible, to enter each of the smaller compartments from the main compartment. In each of the compartments, there were two small (9 cm tall, 7.5 cm wide, 6 cm deep) wooden boxes, with an opening facing the door on the bottom of the cage’s front. The bird had to enter the small compartments to see what is under the box. During the test, there was a small plastic dish under each of the two boxes, in which the mealworm was placed. Pictures taken by EV.

**Figure S2.** Individual reaction norms showing the learning pattern of each of the 17 birds, separated by batches (i.e. birds that were in captivity together), in the perseveration phase (top row) and the new learning phase (bottom row). Batch 1 and 3 are from Site 1, Batch 2 from Site 2, and Batch 4 from Site 3. The shape and colour of the points corresponds to that of the symbol on the rewarding box (both of which alternate with each reversal). The points are scattered horizontally around the integer values to increase visibility. The first point of each individual, at 0 on the X axis, shows their behaviour before the reversal (i.e. their associative learning speed).

**R code:**

i=read.table("reversal learning datafile.txt",header=T)

str(i)

**#Initial choice**

**#Reward position: rewarding vs unrewarding:**

tab1=with(subset(i,i$reversals==0),table(firstsucc))

chisq.test(x=tab1,p=c(0.5,0.5))

**#Lateral side: left vs right**

tab2=with(subset(i,i$reversals==0),table(firstsucc,side))

chisq.test(x=c(tab2[1]+tab2[4],tab2[2]+tab2[3]),p=c(0.5,0.5))

**#Shape: O vs X**

tab3=with(subset(i,i$reversals==0),table(firstsucc,shape))

chisq.test(x=c(tab3[1]+tab3[4],tab3[2]+tab3[3]),p=c(0.5,0.5))

**#Colour: blue vs yellow**

tab4=with(subset(i,i$reversals==0),table(firstsucc,color))

chisq.test(x=c(tab4[1]+tab4[4],tab4[2]+tab4[3]),p=c(0.5,0.5))

**#Perseverance**

**#Model (Table 1A)**

library(MASS)

m1=glmmPQL(firstsucc~reversals+revtotal+sex+age+side+color+shape,

random=~1|site/ringnumber,data=subset(i,i$reversals>0),family=negative.binomial(theta=1))

summary(m1)

var_site1=(1.007518e-05)^2

var_birdID1=(6.274445e-07)^2

var_resid1=(0.4823841)^2

var_total1=var_site1+var_birdID1+var_resid1

var_site1/var_total1

(var_site1+var_birdID1)/var_total1

**#New learning**

**#Model (Table 1B)**

library(MASS)

m2=glmmPQL(learned~reversals+revtotal+sex+age+side+color+shape,

random=~1|site/ringnumber,data=subset(i,i$reversals>0),family=negative.binomial(theta=1))

summary(m2)

var_site2=(0.2130761)^2

var_birdID2=(1.487559e-05)^2

var_resid2=(0.5319185)^2

var_total2=var_site2+var_birdID2+var_resid2

var_site2/var_total2

(var_site2+var_birdID2)/var_total2

**#Main figure (Figure 1) - export as 630 x 890**

library(gplots)

par(mfrow = c(2,1))

par(mar=c(4,4,4,0.5))

par(oma=c(1,1,1,1))

with(subset(i,i$reversals>0),plotmeans(firstsucc~reversals,

xlab="Number of reversals",

ylab="Attempts until first success",

main="A) Perseverance phase",

cex.main = 1.5, cex.lab = 1.2,

ylim=c(0,8),barcol="black"))

with(subset(i,i$reversals>0),plotmeans(learned~reversals,

main="B) New learning phase",

ylab="Attempts until learned",

xlab="Number of reversals",

cex.main = 1.5, cex.lab = 1.2,

ylim = c(0,20),barcol="black"))

**#Supplementary Figure (Figure S2) - export as 1200 x 600**

par(mfrow=c(2,4))

par(mar=c(4,4,0.5,0.5))

par(oma=c(1,1,4,1))

**#Batch 1 - Persistence**

plot(1,xlim = c(-0.1,6.1),ylim=c(0,20), type="n",xlab="",

ylab="Attempts until first success",cex.lab=1.6,cex.axis=1.5,bty="n")

rect(-0.34,-0.8,6.34,20.8,col="gray")

segments(x0=0.7,y0=-0.8,x1=0.7,y1=20.8)

segments(x0=-0.2,y0=i[i$ringnumber=="2KX55067"&i$reversals==0,]$firstsucc,

x1=0.8,y1=i[i$ringnumber=="2KX55067"&i$reversals==1,]$firstsucc,lty=2)

segments(x0=0.8,y0=i[i$ringnumber=="2KX55067"&i$reversals==1,]$firstsucc,

x1=1.8,y1=i[i$ringnumber=="2KX55067"&i$reversals==2,]$firstsucc,lty=2)

segments(x0=1.8,y0=i[i$ringnumber=="2KX55067"&i$reversals==2,]$firstsucc,

x1=2.8,y1=i[i$ringnumber=="2KX55067"&i$reversals==3,]$firstsucc,lty=2)

segments(x0=2.8,y0=i[i$ringnumber=="2KX55067"&i$reversals==3,]$firstsucc,

x1=3.8,y1=i[i$ringnumber=="2KX55067"&i$reversals==4,]$firstsucc,lty=2)

with(subset(i,i$ringnumber=="2KX55067"),

points(x=reversals-0.2,y=firstsucc,pch=c(1,4,1,4,1),col=as.character(color),cex=1.4))

segments(x0=-0.1,y0=i[i$ringnumber=="2KX68977"&i$reversals==0,]$firstsucc,

x1=0.9,y1=i[i$ringnumber=="2KX68977"&i$reversals==1,]$firstsucc,lty=2)

segments(x0=0.9,y0=i[i$ringnumber=="2KX68977"&i$reversals==1,]$firstsucc,

x1=1.9,y1=i[i$ringnumber=="2KX68977"&i$reversals==2,]$firstsucc,lty=2)

segments(x0=1.9,y0=i[i$ringnumber=="2KX68977"&i$reversals==2,]$firstsucc,

x1=2.9,y1=i[i$ringnumber=="2KX68977"&i$reversals==3,]$firstsucc,lty=2)

segments(x0=2.9,y0=i[i$ringnumber=="2KX68977"&i$reversals==3,]$firstsucc,

x1=3.9,y1=i[i$ringnumber=="2KX68977"&i$reversals==4,]$firstsucc,lty=2)

with(subset(i,i$ringnumber=="2KX68977"),

points(x=reversals-0.1,y=firstsucc,pch=c(4,1,4,1,4),col=as.character(color),cex=1.4))

segments(x0=0,y0=i[i$ringnumber=="2KZ09018"&i$reversals==0,]$firstsucc,

x1=1,y1=i[i$ringnumber=="2KZ09018"&i$reversals==1,]$firstsucc,lty=2)

segments(x0=1,y0=i[i$ringnumber=="2KZ09018"&i$reversals==1,]$firstsucc,

x1=2,y1=i[i$ringnumber=="2KZ09018"&i$reversals==2,]$firstsucc,lty=2)

segments(x0=2,y0=i[i$ringnumber=="2KZ09018"&i$reversals==2,]$firstsucc,

x1=3,y1=i[i$ringnumber=="2KZ09018"&i$reversals==3,]$firstsucc,lty=2)

segments(x0=3,y0=i[i$ringnumber=="2KZ09018"&i$reversals==3,]$firstsucc,

x1=4,y1=i[i$ringnumber=="2KZ09018"&i$reversals==4,]$firstsucc,lty=2)

with(subset(i,i$ringnumber=="2KZ09018"),

points(x=reversals,y=firstsucc,pch=c(1,4,1,4,1),col=as.character(color),cex=1.4))

segments(x0=0.1,y0=i[i$ringnumber=="2KZ09019"&i$reversals==0,]$firstsucc,

x1=1.1,y1=i[i$ringnumber=="2KZ09019"&i$reversals==1,]$firstsucc,lty=2)

segments(x0=1.1,y0=i[i$ringnumber=="2KZ09019"&i$reversals==1,]$firstsucc,

x1=2.1,y1=i[i$ringnumber=="2KZ09019"&i$reversals==2,]$firstsucc,lty=2)

segments(x0=2.1,y0=i[i$ringnumber=="2KZ09019"&i$reversals==2,]$firstsucc,

x1=3.1,y1=i[i$ringnumber=="2KZ09019"&i$reversals==3,]$firstsucc,lty=2)

segments(x0=3.1,y0=i[i$ringnumber=="2KZ09019"&i$reversals==3,]$firstsucc,

x1=4.1,y1=i[i$ringnumber=="2KZ09019"&i$reversals==4,]$firstsucc,lty=2)

segments(x0=4.1,y0=i[i$ringnumber=="2KZ09019"&i$reversals==4,]$firstsucc,

x1=5.1,y1=i[i$ringnumber=="2KZ09019"&i$reversals==5,]$firstsucc,lty=2)

segments(x0=5.1,y0=i[i$ringnumber=="2KZ09019"&i$reversals==5,]$firstsucc,

x1=6.1,y1=i[i$ringnumber=="2KZ09019"&i$reversals==6,]$firstsucc,lty=2)

with(subset(i,i$ringnumber=="2KZ09019"),

points(x=reversals+0.1,y=firstsucc,pch=c(4,1,4,1,4,1,4),col=as.character(color),cex=1.4))

segments(x0=0.2,y0=i[i$ringnumber=="2KZ09020"&i$reversals==0,]$firstsucc,

x1=1.2,y1=i[i$ringnumber=="2KZ09020"&i$reversals==1,]$firstsucc,lty=2)

segments(x0=1.2,y0=i[i$ringnumber=="2KZ09020"&i$reversals==1,]$firstsucc,

x1=2.2,y1=i[i$ringnumber=="2KZ09020"&i$reversals==2,]$firstsucc,lty=2)

segments(x0=2.2,y0=i[i$ringnumber=="2KZ09020"&i$reversals==2,]$firstsucc,

x1=3.2,y1=i[i$ringnumber=="2KZ09020"&i$reversals==3,]$firstsucc,lty=2)

segments(x0=3.2,y0=i[i$ringnumber=="2KZ09020"&i$reversals==3,]$firstsucc,

x1=4.2,y1=i[i$ringnumber=="2KZ09020"&i$reversals==4,]$firstsucc,lty=2)

segments(x0=4.2,y0=i[i$ringnumber=="2KZ09020"&i$reversals==4,]$firstsucc,

x1=5.2,y1=i[i$ringnumber=="2KZ09020"&i$reversals==5,]$firstsucc,lty=2)

with(subset(i,i$ringnumber=="2KZ09020"),

points(x=reversals+0.2,y=firstsucc,pch=c(1,4,1,4,1,4),col=as.character(color),cex=1.4))

**#Batch 2 - Persistence**

plot(1,xlim = c(-0.1,6.1),ylim=c(0,20), type="n",xlab="",ylab="",cex.lab=1.6,cex.axis=1.5,bty="n")

rect(-0.34,-0.8,6.34,20.8,col="gray")

segments(x0=0.7,y0=-0.8,x1=0.7,y1=20.8)

segments(x0=0,y0=i[i$ringnumber=="2KZ09021"&i$reversals==0,]$firstsucc,

x1=1,y1=i[i$ringnumber=="2KZ09021"&i$reversals==1,]$firstsucc,lty=2)

segments(x0=1,y0=i[i$ringnumber=="2KZ09021"&i$reversals==1,]$firstsucc,

x1=2,y1=i[i$ringnumber=="2KZ09021"&i$reversals==2,]$firstsucc,lty=2)

segments(x0=2,y0=i[i$ringnumber=="2KZ09021"&i$reversals==2,]$firstsucc,

x1=3,y1=i[i$ringnumber=="2KZ09021"&i$reversals==3,]$firstsucc,lty=2)

segments(x0=3,y0=i[i$ringnumber=="2KZ09021"&i$reversals==3,]$firstsucc,

x1=4,y1=i[i$ringnumber=="2KZ09021"&i$reversals==4,]$firstsucc,lty=2)

segments(x0=4,y0=i[i$ringnumber=="2KZ09021"&i$reversals==4,]$firstsucc,

x1=5,y1=i[i$ringnumber=="2KZ09021"&i$reversals==5,]$firstsucc,lty=2)

segments(x0=5,y0=i[i$ringnumber=="2KZ09021"&i$reversals==5,]$firstsucc,

x1=6,y1=i[i$ringnumber=="2KZ09021"&i$reversals==6,]$firstsucc,lty=2)

with(subset(i,i$ringnumber=="2KZ09021"),

points(x=reversals,y=firstsucc,pch=c(1,4,1,4,1,4,1),col=as.character(color),cex=1.4))

segments(x0=-0.2,y0=i[i$ringnumber=="2KZ09023"&i$reversals==0,]$firstsucc,

x1=0.8,y1=i[i$ringnumber=="2KZ09023"&i$reversals==1,]$firstsucc,lty=2)

segments(x0=0.8,y0=i[i$ringnumber=="2KZ09023"&i$reversals==1,]$firstsucc,

x1=1.8,y1=i[i$ringnumber=="2KZ09023"&i$reversals==2,]$firstsucc,lty=2)

segments(x0=1.8,y0=i[i$ringnumber=="2KZ09023"&i$reversals==2,]$firstsucc,

x1=2.8,y1=i[i$ringnumber=="2KZ09023"&i$reversals==3,]$firstsucc,lty=2)

with(subset(i,i$ringnumber=="2KZ09023"),

points(x=reversals-0.2,y=firstsucc,pch=c(1,4,1,4),col=as.character(color),cex=1.4))

segments(x0=-0.1,y0=i[i$ringnumber=="2KZ09024"&i$reversals==0,]$firstsucc,

x1=0.9,y1=i[i$ringnumber=="2KZ09024"&i$reversals==1,]$firstsucc,lty=2)

segments(x0=0.9,y0=i[i$ringnumber=="2KZ09024"&i$reversals==1,]$firstsucc,

x1=1.9,y1=i[i$ringnumber=="2KZ09024"&i$reversals==2,]$firstsucc,lty=2)

segments(x0=1.9,y0=i[i$ringnumber=="2KZ09024"&i$reversals==2,]$firstsucc,

x1=2.9,y1=i[i$ringnumber=="2KZ09024"&i$reversals==3,]$firstsucc,lty=2)

segments(x0=2.9,y0=i[i$ringnumber=="2KZ09024"&i$reversals==3,]$firstsucc,

x1=3.9,y1=i[i$ringnumber=="2KZ09024"&i$reversals==4,]$firstsucc,lty=2)

segments(x0=3.9,y0=i[i$ringnumber=="2KZ09024"&i$reversals==4,]$firstsucc,

x1=4.9,y1=i[i$ringnumber=="2KZ09024"&i$reversals==5,]$firstsucc,lty=2)

segments(x0=4.9,y0=i[i$ringnumber=="2KZ09024"&i$reversals==5,]$firstsucc,

x1=5.9,y1=i[i$ringnumber=="2KZ09024"&i$reversals==6,]$firstsucc,lty=2)

with(subset(i,i$ringnumber=="2KZ09024"),

points(x=reversals-0.1,y=firstsucc,pch=c(4,1,4,1,4,1,4),col=as.character(color),cex=1.4))

segments(x0=0.1,y0=i[i$ringnumber=="2KZ09025"&i$reversals==0,]$firstsucc,

x1=1.1,y1=i[i$ringnumber=="2KZ09025"&i$reversals==1,]$firstsucc,lty=2)

segments(x0=1.1,y0=i[i$ringnumber=="2KZ09025"&i$reversals==1,]$firstsucc,

x1=2.1,y1=i[i$ringnumber=="2KZ09025"&i$reversals==2,]$firstsucc,lty=2)

segments(x0=2.1,y0=i[i$ringnumber=="2KZ09025"&i$reversals==2,]$firstsucc,

x1=3.1,y1=i[i$ringnumber=="2KZ09025"&i$reversals==3,]$firstsucc,lty=2)

segments(x0=3.1,y0=i[i$ringnumber=="2KZ09025"&i$reversals==3,]$firstsucc,

x1=4.1,y1=i[i$ringnumber=="2KZ09025"&i$reversals==4,]$firstsucc,lty=2)

segments(x0=4.1,y0=i[i$ringnumber=="2KZ09025"&i$reversals==4,]$firstsucc,

x1=5.1,y1=i[i$ringnumber=="2KZ09025"&i$reversals==5,]$firstsucc,lty=2)

segments(x0=5.1,y0=i[i$ringnumber=="2KZ09025"&i$reversals==5,]$firstsucc,

x1=6.1,y1=i[i$ringnumber=="2KZ09025"&i$reversals==6,]$firstsucc,lty=2)

with(subset(i,i$ringnumber=="2KZ09025"),

points(x=reversals+0.1,y=firstsucc,pch=c(4,1,4,1,4,1,4),col=as.character(color,cex=1.4)))

**#Batch 3 - Persistence**

plot(1,xlim = c(-0.1,6.1),ylim=c(0,20), type="n",xlab="",ylab="",cex.lab=1.6,cex.axis=1.5,bty="n")

rect(-0.34,-0.8,6.34,20.8,col="gray")

segments(x0=0.7,y0=-0.8,x1=0.7,y1=20.8)

segments(x0=0,y0=i[i$ringnumber=="2KZ09027"&i$reversals==0,]$firstsucc,

x1=1,y1=i[i$ringnumber=="2KZ09027"&i$reversals==1,]$firstsucc,lty=2)

segments(x0=1,y0=i[i$ringnumber=="2KZ09027"&i$reversals==1,]$firstsucc,

x1=2,y1=i[i$ringnumber=="2KZ09027"&i$reversals==2,]$firstsucc,lty=2)

segments(x0=2,y0=i[i$ringnumber=="2KZ09027"&i$reversals==2,]$firstsucc,

x1=3,y1=i[i$ringnumber=="2KZ09027"&i$reversals==3,]$firstsucc,lty=2)

with(subset(i,i$ringnumber=="2KZ09027"),

points(x=reversals,y=firstsucc,pch=c(1,4,1,4),col=as.character(color),cex=1.4))

segments(x0=0.1,y0=i[i$ringnumber=="2KZ09028"&i$reversals==0,]$firstsucc,

x1=1.1,y1=i[i$ringnumber=="2KZ09028"&i$reversals==1,]$firstsucc,lty=2)

segments(x0=1.1,y0=i[i$ringnumber=="2KZ09028"&i$reversals==1,]$firstsucc,

x1=2.1,y1=i[i$ringnumber=="2KZ09028"&i$reversals==2,]$firstsucc,lty=2)

with(subset(i,i$ringnumber=="2KZ09028"),

points(x=reversals+0.1,y=firstsucc,pch=c(1,4,1),col=as.character(color),cex=1.4))

segments(x0=-0.1,y0=i[i$ringnumber=="2KZ09029"&i$reversals==0,]$firstsucc,

x1=0.9,y1=i[i$ringnumber=="2KZ09029"&i$reversals==1,]$firstsucc,lty=2)

segments(x0=0.9,y0=i[i$ringnumber=="2KZ09029"&i$reversals==1,]$firstsucc,

x1=1.9,y1=i[i$ringnumber=="2KZ09029"&i$reversals==2,]$firstsucc,lty=2)

segments(x0=1.9,y0=i[i$ringnumber=="2KZ09029"&i$reversals==2,]$firstsucc,

x1=2.9,y1=i[i$ringnumber=="2KZ09029"&i$reversals==3,]$firstsucc,lty=2)

segments(x0=2.9,y0=i[i$ringnumber=="2KZ09029"&i$reversals==3,]$firstsucc,

x1=3.9,y1=i[i$ringnumber=="2KZ09029"&i$reversals==4,]$firstsucc,lty=2)

with(subset(i,i$ringnumber=="2KZ09029"),

points(x=reversals-0.1,y=firstsucc,pch=c(1,4,1,4,1),col=as.character(color),cex=1.4))

**#Batch 4 - Persistence**

plot(1,xlim = c(-0.1,6.1),ylim=c(0,20), type="n",xlab="",ylab="",cex.lab=1.6,cex.axis=1.5,bty="n")

rect(-0.34,-0.8,6.34,20.8,col="gray")

segments(x0=0.7,y0=-0.8,x1=0.7,y1=20.8)

segments(x0=0,y0=i[i$ringnumber=="2KZ09030"&i$reversals==0,]$firstsucc,

x1=1,y1=i[i$ringnumber=="2KZ09030"&i$reversals==1,]$firstsucc,lty=2)

segments(x0=1,y0=i[i$ringnumber=="2KZ09030"&i$reversals==1,]$firstsucc,

x1=2,y1=i[i$ringnumber=="2KZ09030"&i$reversals==2,]$firstsucc,lty=2)

segments(x0=2,y0=i[i$ringnumber=="2KZ09030"&i$reversals==2,]$firstsucc,

x1=3,y1=i[i$ringnumber=="2KZ09030"&i$reversals==3,]$firstsucc,lty=2)

segments(x0=3,y0=i[i$ringnumber=="2KZ09030"&i$reversals==3,]$firstsucc,

x1=4,y1=i[i$ringnumber=="2KZ09030"&i$reversals==4,]$firstsucc,lty=2)

segments(x0=4,y0=i[i$ringnumber=="2KZ09030"&i$reversals==4,]$firstsucc,

x1=5,y1=i[i$ringnumber=="2KZ09030"&i$reversals==5,]$firstsucc,lty=2)

with(subset(i,i$ringnumber=="2KZ09030"),

points(x=reversals,y=firstsucc,pch=c(4,1,4,1,4,1),col=as.character(color),cex=1.4))

segments(x0=0.2,y0=i[i$ringnumber=="2KZ09031"&i$reversals==0,]$firstsucc,

x1=1.2,y1=i[i$ringnumber=="2KZ09031"&i$reversals==1,]$firstsucc,lty=2)

segments(x0=1.2,y0=i[i$ringnumber=="2KZ09031"&i$reversals==1,]$firstsucc,

x1=2.2,y1=i[i$ringnumber=="2KZ09031"&i$reversals==2,]$firstsucc,lty=2)

segments(x0=2.2,y0=i[i$ringnumber=="2KZ09031"&i$reversals==2,]$firstsucc,

x1=3.2,y1=i[i$ringnumber=="2KZ09031"&i$reversals==3,]$firstsucc,lty=2)

segments(x0=3.2,y0=i[i$ringnumber=="2KZ09031"&i$reversals==3,]$firstsucc,

x1=4.2,y1=i[i$ringnumber=="2KZ09031"&i$reversals==4,]$firstsucc,lty=2)

with(subset(i,i$ringnumber=="2KZ09031"),

points(x=reversals+0.2,y=firstsucc,pch=c(4,1,4,1,4),col=as.character(color),cex=1.4))

segments(x0=0.1,y0=i[i$ringnumber=="2KZ09032"&i$reversals==0,]$firstsucc,

x1=1.1,y1=i[i$ringnumber=="2KZ09032"&i$reversals==1,]$firstsucc,lty=2)

segments(x0=1.1,y0=i[i$ringnumber=="2KZ09032"&i$reversals==1,]$firstsucc,

x1=2.1,y1=i[i$ringnumber=="2KZ09032"&i$reversals==2,]$firstsucc,lty=2)

segments(x0=2.1,y0=i[i$ringnumber=="2KZ09032"&i$reversals==2,]$firstsucc,

x1=3.1,y1=i[i$ringnumber=="2KZ09032"&i$reversals==3,]$firstsucc,lty=2)

segments(x0=3.1,y0=i[i$ringnumber=="2KZ09032"&i$reversals==3,]$firstsucc,

x1=4.1,y1=i[i$ringnumber=="2KZ09032"&i$reversals==4,]$firstsucc,lty=2)

with(subset(i,i$ringnumber=="2KZ09032"),

points(x=reversals+0.1,y=firstsucc,pch=c(4,1,4,1,4),col=as.character(color),cex=1.4))

segments(x0=-0.1,y0=i[i$ringnumber=="2KZ09033"&i$reversals==0,]$firstsucc,

x1=0.9,y1=i[i$ringnumber=="2KZ09033"&i$reversals==1,]$firstsucc,lty=2)

segments(x0=0.9,y0=i[i$ringnumber=="2KZ09033"&i$reversals==1,]$firstsucc,

x1=1.9,y1=i[i$ringnumber=="2KZ09033"&i$reversals==2,]$firstsucc,lty=2)

segments(x0=1.9,y0=i[i$ringnumber=="2KZ09033"&i$reversals==2,]$firstsucc,

x1=2.9,y1=i[i$ringnumber=="2KZ09033"&i$reversals==3,]$firstsucc,lty=2)

segments(x0=2.9,y0=i[i$ringnumber=="2KZ09033"&i$reversals==3,]$firstsucc,

x1=3.9,y1=i[i$ringnumber=="2KZ09033"&i$reversals==4,]$firstsucc,lty=2)

with(subset(i,i$ringnumber=="2KZ09033"),

points(x=reversals-0.1,y=firstsucc,pch=c(1,4,1,4,1),col=as.character(color),cex=1.4))

segments(x0=-0.2,y0=i[i$ringnumber=="2KZ09034"&i$reversals==0,]$firstsucc,

x1=0.8,y1=i[i$ringnumber=="2KZ09034"&i$reversals==1,]$firstsucc,lty=2)

segments(x0=0.8,y0=i[i$ringnumber=="2KZ09034"&i$reversals==1,]$firstsucc,

x1=1.8,y1=i[i$ringnumber=="2KZ09034"&i$reversals==2,]$firstsucc,lty=2)

segments(x0=1.8,y0=i[i$ringnumber=="2KZ09034"&i$reversals==2,]$firstsucc,

x1=2.8,y1=i[i$ringnumber=="2KZ09034"&i$reversals==3,]$firstsucc,lty=2)

segments(x0=2.8,y0=i[i$ringnumber=="2KZ09034"&i$reversals==3,]$firstsucc,

x1=3.8,y1=i[i$ringnumber=="2KZ09034"&i$reversals==4,]$firstsucc,lty=2)

segments(x0=3.8,y0=i[i$ringnumber=="2KZ09034"&i$reversals==4,]$firstsucc,

x1=4.8,y1=i[i$ringnumber=="2KZ09034"&i$reversals==5,]$firstsucc,lty=2)

segments(x0=4.8,y0=i[i$ringnumber=="2KZ09034"&i$reversals==5,]$firstsucc,

x1=5.8,y1=i[i$ringnumber=="2KZ09034"&i$reversals==6,]$firstsucc,lty=2)

with(subset(i,i$ringnumber=="2KZ09034"),

points(x=reversals-0.2,y=firstsucc,pch=c(4,1,4,1,4,1,4),col=as.character(color),cex=1.4))

**#Batch 1 - New learning**

plot(1,xlim = c(-0.1,6.1),ylim=c(0,35), type="n",xlab="Number of reversals",

ylab="Attempts until learned",cex.lab=1.6,cex.axis=1.5,bty="n")

rect(-0.34,-1.4,6.34,36.4,col="gray")

segments(x0=0.7,y0=-1.4,x1=0.7,y1=36.4)

segments(x0=-0.2,y0=i[i$ringnumber=="2KX55067"&i$reversals==0,]$learned,

x1=0.8,y1=i[i$ringnumber=="2KX55067"&i$reversals==1,]$learned,lty=2)

segments(x0=0.8,y0=i[i$ringnumber=="2KX55067"&i$reversals==1,]$learned,

x1=1.8,y1=i[i$ringnumber=="2KX55067"&i$reversals==2,]$learned,lty=2)

segments(x0=1.8,y0=i[i$ringnumber=="2KX55067"&i$reversals==2,]$learned,

x1=2.8,y1=i[i$ringnumber=="2KX55067"&i$reversals==3,]$learned,lty=2)

with(subset(i,i$ringnumber=="2KX55067"),

points(x=reversals-0.2,y=learned,pch=c(1,4,1,4),col=as.character(color),cex=1.4))

segments(x0=-0.1,y0=i[i$ringnumber=="2KX68977"&i$reversals==0,]$learned,

x1=0.9,y1=i[i$ringnumber=="2KX68977"&i$reversals==1,]$learned,lty=2)

segments(x0=0.9,y0=i[i$ringnumber=="2KX68977"&i$reversals==1,]$learned,

x1=1.9,y1=i[i$ringnumber=="2KX68977"&i$reversals==2,]$learned,lty=2)

segments(x0=1.9,y0=i[i$ringnumber=="2KX68977"&i$reversals==2,]$learned,

x1=2.9,y1=i[i$ringnumber=="2KX68977"&i$reversals==3,]$learned,lty=2)

with(subset(i,i$ringnumber=="2KX68977"),

points(x=reversals-0.1,y=learned,pch=c(4,1,4,1),col=as.character(color),cex=1.4))

segments(x0=0,y0=i[i$ringnumber=="2KZ09018"&i$reversals==0,]$learned,

x1=1,y1=i[i$ringnumber=="2KZ09018"&i$reversals==1,]$learned,lty=2)

segments(x0=1,y0=i[i$ringnumber=="2KZ09018"&i$reversals==1,]$learned,

x1=2,y1=i[i$ringnumber=="2KZ09018"&i$reversals==2,]$learned,lty=2)

segments(x0=2,y0=i[i$ringnumber=="2KZ09018"&i$reversals==2,]$learned,

x1=3,y1=i[i$ringnumber=="2KZ09018"&i$reversals==3,]$learned,lty=2)

with(subset(i,i$ringnumber=="2KZ09018"),

points(x=reversals-0.08,y=learned,pch=c(1,4,1,4),col=as.character(color),cex=1.4))

segments(x0=0.1,y0=i[i$ringnumber=="2KZ09019"&i$reversals==0,]$learned,

x1=1.1,y1=i[i$ringnumber=="2KZ09019"&i$reversals==1,]$learned,lty=2)

segments(x0=1.1,y0=i[i$ringnumber=="2KZ09019"&i$reversals==1,]$learned,

x1=2.1,y1=i[i$ringnumber=="2KZ09019"&i$reversals==2,]$learned,lty=2)

segments(x0=2.1,y0=i[i$ringnumber=="2KZ09019"&i$reversals==2,]$learned,

x1=3.1,y1=i[i$ringnumber=="2KZ09019"&i$reversals==3,]$learned,lty=2)

segments(x0=3.1,y0=i[i$ringnumber=="2KZ09019"&i$reversals==3,]$learned,

x1=4.1,y1=i[i$ringnumber=="2KZ09019"&i$reversals==4,]$learned,lty=2)

segments(x0=4.1,y0=i[i$ringnumber=="2KZ09019"&i$reversals==4,]$learned,

x1=5.1,y1=i[i$ringnumber=="2KZ09019"&i$reversals==5,]$learned,lty=2)

with(subset(i,i$ringnumber=="2KZ09019"),

points(x=reversals+0.1,y=learned,pch=c(4,1,4,1,4,1),col=as.character(color),cex=1.4))

segments(x0=0.2,y0=i[i$ringnumber=="2KZ09020"&i$reversals==0,]$learned,

x1=1.2,y1=i[i$ringnumber=="2KZ09020"&i$reversals==1,]$learned,lty=2)

segments(x0=1.2,y0=i[i$ringnumber=="2KZ09020"&i$reversals==1,]$learned,

x1=2.2,y1=i[i$ringnumber=="2KZ09020"&i$reversals==2,]$learned,lty=2)

segments(x0=2.2,y0=i[i$ringnumber=="2KZ09020"&i$reversals==2,]$learned,

x1=3.2,y1=i[i$ringnumber=="2KZ09020"&i$reversals==3,]$learned,lty=2)

segments(x0=3.2,y0=i[i$ringnumber=="2KZ09020"&i$reversals==3,]$learned,

x1=4.2,y1=i[i$ringnumber=="2KZ09020"&i$reversals==4,]$learned,lty=2)

with(subset(i,i$ringnumber=="2KZ09020"),

points(x=reversals+0.2,y=learned,pch=c(1,4,1,4,1),col=as.character(color),cex=1.4))

**#Batch 2 - New learning**

plot(1,xlim = c(-0.1,6.1),ylim=c(0,35), type="n",xlab="Number of reversals",ylab="",cex.lab=1.6,cex.axis=1.5,bty="n")

rect(-0.34,-1.4,6.34,36.4,col="gray")

segments(x0=0.7,y0=-1.4,x1=0.7,y1=36.4)

segments(x0=0,y0=i[i$ringnumber=="2KZ09021"&i$reversals==0,]$learned,

x1=1,y1=i[i$ringnumber=="2KZ09021"&i$reversals==1,]$learned,lty=2)

segments(x0=1,y0=i[i$ringnumber=="2KZ09021"&i$reversals==1,]$learned,

x1=2,y1=i[i$ringnumber=="2KZ09021"&i$reversals==2,]$learned,lty=2)

segments(x0=2,y0=i[i$ringnumber=="2KZ09021"&i$reversals==2,]$learned,

x1=3,y1=i[i$ringnumber=="2KZ09021"&i$reversals==3,]$learned,lty=2)

segments(x0=3,y0=i[i$ringnumber=="2KZ09021"&i$reversals==3,]$learned,

x1=4,y1=i[i$ringnumber=="2KZ09021"&i$reversals==4,]$learned,lty=2)

segments(x0=4,y0=i[i$ringnumber=="2KZ09021"&i$reversals==4,]$learned,

x1=5,y1=i[i$ringnumber=="2KZ09021"&i$reversals==5,]$learned,lty=2)

with(subset(i,i$ringnumber=="2KZ09021"),

points(x=reversals,y=learned,pch=c(1,4,1,4,1,4),col=as.character(color),cex=1.4))

segments(x0=-0.2,y0=i[i$ringnumber=="2KZ09023"&i$reversals==0,]$learned,

x1=0.8,y1=i[i$ringnumber=="2KZ09023"&i$reversals==1,]$learned,lty=2)

segments(x0=0.8,y0=i[i$ringnumber=="2KZ09023"&i$reversals==1,]$learned,

x1=1.8,y1=i[i$ringnumber=="2KZ09023"&i$reversals==2,]$learned,lty=2)

with(subset(i,i$ringnumber=="2KZ09023"),

points(x=reversals-0.2,y=learned,pch=c(1,4,1),col=as.character(color),cex=1.4))

segments(x0=-0.1,y0=i[i$ringnumber=="2KZ09024"&i$reversals==0,]$learned,

x1=0.9,y1=i[i$ringnumber=="2KZ09024"&i$reversals==1,]$learned,lty=2)

segments(x0=0.9,y0=i[i$ringnumber=="2KZ09024"&i$reversals==1,]$learned,

x1=1.9,y1=i[i$ringnumber=="2KZ09024"&i$reversals==2,]$learned,lty=2)

segments(x0=1.9,y0=i[i$ringnumber=="2KZ09024"&i$reversals==2,]$learned,

x1=2.9,y1=i[i$ringnumber=="2KZ09024"&i$reversals==3,]$learned,lty=2)

segments(x0=2.9,y0=i[i$ringnumber=="2KZ09024"&i$reversals==3,]$learned,

x1=3.9,y1=i[i$ringnumber=="2KZ09024"&i$reversals==4,]$learned,lty=2)

segments(x0=3.9,y0=i[i$ringnumber=="2KZ09024"&i$reversals==4,]$learned,

x1=4.9,y1=i[i$ringnumber=="2KZ09024"&i$reversals==5,]$learned,lty=2)

with(subset(i,i$ringnumber=="2KZ09024"),

points(x=reversals-0.1,y=learned,pch=c(4,1,4,1,4,1),col=as.character(color),cex=1.4))

segments(x0=0.1,y0=i[i$ringnumber=="2KZ09025"&i$reversals==0,]$learned,

x1=1.1,y1=i[i$ringnumber=="2KZ09025"&i$reversals==1,]$learned,lty=2)

segments(x0=1.1,y0=i[i$ringnumber=="2KZ09025"&i$reversals==1,]$learned,

x1=2.1,y1=i[i$ringnumber=="2KZ09025"&i$reversals==2,]$learned,lty=2)

segments(x0=2.1,y0=i[i$ringnumber=="2KZ09025"&i$reversals==2,]$learned,

x1=3.1,y1=i[i$ringnumber=="2KZ09025"&i$reversals==3,]$learned,lty=2)

segments(x0=3.1,y0=i[i$ringnumber=="2KZ09025"&i$reversals==3,]$learned,

x1=4.1,y1=i[i$ringnumber=="2KZ09025"&i$reversals==4,]$learned,lty=2)

segments(x0=4.1,y0=i[i$ringnumber=="2KZ09025"&i$reversals==4,]$learned,

x1=5.1,y1=i[i$ringnumber=="2KZ09025"&i$reversals==5,]$learned,lty=2)

with(subset(i,i$ringnumber=="2KZ09025"),

points(x=reversals+0.1,y=learned,pch=c(4,1,4,1,4,1),col=as.character(color),cex=1.4))

**#Batch 3 - New learning**

plot(1,xlim = c(-0.1,6.1),ylim=c(0,35), type="n",xlab="Number of reversals",ylab="",cex.lab=1.6,cex.axis=1.5,bty="n")

rect(-0.34,-1.4,6.34,36.4,col="gray")

segments(x0=0.7,y0=-1.4,x1=0.7,y1=36.4)

segments(x0=0,y0=i[i$ringnumber=="2KZ09027"&i$reversals==0,]$learned,

x1=1,y1=i[i$ringnumber=="2KZ09027"&i$reversals==1,]$learned,lty=2)

segments(x0=1,y0=i[i$ringnumber=="2KZ09027"&i$reversals==1,]$learned,

x1=2,y1=i[i$ringnumber=="2KZ09027"&i$reversals==2,]$learned,lty=2)

with(subset(i,i$ringnumber=="2KZ09027"),

points(x=reversals,y=learned,pch=c(1,4,1),col=as.character(color),cex=1.4))

segments(x0=0.1,y0=i[i$ringnumber=="2KZ09028"&i$reversals==0,]$learned,

x1=1.1,y1=i[i$ringnumber=="2KZ09028"&i$reversals==1,]$learned,lty=2)

segments(x0=1.1,y0=i[i$ringnumber=="2KZ09028"&i$reversals==1,]$learned,

x1=2.1,y1=i[i$ringnumber=="2KZ09028"&i$reversals==2,]$learned,lty=2)

with(subset(i,i$ringnumber=="2KZ09028"),

points(x=reversals+0.1,y=learned,pch=c(1,4,1),col=as.character(color),cex=1.4))

segments(x0=-0.1,y0=i[i$ringnumber=="2KZ09029"&i$reversals==0,]$learned,

x1=0.9,y1=i[i$ringnumber=="2KZ09029"&i$reversals==1,]$learned,lty=2)

segments(x0=0.9,y0=i[i$ringnumber=="2KZ09029"&i$reversals==1,]$learned,

x1=1.9,y1=i[i$ringnumber=="2KZ09029"&i$reversals==2,]$learned,lty=2)

segments(x0=1.9,y0=i[i$ringnumber=="2KZ09029"&i$reversals==2,]$learned,

x1=2.9,y1=i[i$ringnumber=="2KZ09029"&i$reversals==3,]$learned,lty=2)

with(subset(i,i$ringnumber=="2KZ09029"),

points(x=reversals-0.1,y=learned,pch=c(1,4,1,4),col=as.character(color),cex=1.4))

**#Batch 4 - New learning**

plot(1,xlim = c(-0.1,6.1),ylim=c(0,35), type="n",xlab="Number of reversals",ylab="",cex.lab=1.6,cex.axis=1.5,bty="n")

rect(-0.34,-1.4,6.34,36.4,col="gray")

segments(x0=0.7,y0=-1.4,x1=0.7,y1=36.4)

segments(x0=0,y0=i[i$ringnumber=="2KZ09030"&i$reversals==0,]$learned,

x1=1,y1=i[i$ringnumber=="2KZ09030"&i$reversals==1,]$learned,lty=2)

segments(x0=1,y0=i[i$ringnumber=="2KZ09030"&i$reversals==1,]$learned,

x1=2,y1=i[i$ringnumber=="2KZ09030"&i$reversals==2,]$learned,lty=2)

segments(x0=2,y0=i[i$ringnumber=="2KZ09030"&i$reversals==2,]$learned,

x1=3,y1=i[i$ringnumber=="2KZ09030"&i$reversals==3,]$learned,lty=2)

segments(x0=3,y0=i[i$ringnumber=="2KZ09030"&i$reversals==3,]$learned,

x1=4,y1=i[i$ringnumber=="2KZ09030"&i$reversals==4,]$learned,lty=2)

with(subset(i,i$ringnumber=="2KZ09030"),

points(x=reversals,y=learned,pch=c(4,1,4,1,4),col=as.character(color),cex=1.4))

segments(x0=0.2,y0=i[i$ringnumber=="2KZ09031"&i$reversals==0,]$learned,

x1=1.2,y1=i[i$ringnumber=="2KZ09031"&i$reversals==1,]$learned,lty=2)

segments(x0=1.2,y0=i[i$ringnumber=="2KZ09031"&i$reversals==1,]$learned,

x1=2.2,y1=i[i$ringnumber=="2KZ09031"&i$reversals==2,]$learned,lty=2)

segments(x0=2.2,y0=i[i$ringnumber=="2KZ09031"&i$reversals==2,]$learned,

x1=3.2,y1=i[i$ringnumber=="2KZ09031"&i$reversals==3,]$learned,lty=2)

with(subset(i,i$ringnumber=="2KZ09031"),

points(x=reversals+0.2,y=learned,pch=c(4,1,4,1),col=as.character(color),cex=1.4))

segments(x0=0.1,y0=i[i$ringnumber=="2KZ09032"&i$reversals==0,]$learned,

x1=1.1,y1=i[i$ringnumber=="2KZ09032"&i$reversals==1,]$learned,lty=2)

segments(x0=1.1,y0=i[i$ringnumber=="2KZ09032"&i$reversals==1,]$learned,

x1=2.1,y1=i[i$ringnumber=="2KZ09032"&i$reversals==2,]$learned,lty=2)

segments(x0=2.1,y0=i[i$ringnumber=="2KZ09032"&i$reversals==2,]$learned,

x1=3.1,y1=i[i$ringnumber=="2KZ09032"&i$reversals==3,]$learned,lty=2)

with(subset(i,i$ringnumber=="2KZ09032"),

points(x=reversals+0.1,y=learned,pch=c(4,1,4,1),col=as.character(color),cex=1.4))

segments(x0=-0.1,y0=i[i$ringnumber=="2KZ09033"&i$reversals==0,]$learned,

x1=0.9,y1=i[i$ringnumber=="2KZ09033"&i$reversals==1,]$learned,lty=2)

segments(x0=0.9,y0=i[i$ringnumber=="2KZ09033"&i$reversals==1,]$learned,

x1=1.9,y1=i[i$ringnumber=="2KZ09033"&i$reversals==2,]$learned,lty=2)

segments(x0=1.9,y0=i[i$ringnumber=="2KZ09033"&i$reversals==2,]$learned,

x1=2.9,y1=i[i$ringnumber=="2KZ09033"&i$reversals==3,]$learned,lty=2)

with(subset(i,i$ringnumber=="2KZ09033"),

points(x=reversals-0.1,y=learned,pch=c(1,4,1,4),col=as.character(color),cex=1.4))

segments(x0=-0.2,y0=i[i$ringnumber=="2KZ09034"&i$reversals==0,]$learned,

x1=0.8,y1=i[i$ringnumber=="2KZ09034"&i$reversals==1,]$learned,lty=2)

segments(x0=0.8,y0=i[i$ringnumber=="2KZ09034"&i$reversals==1,]$learned,

x1=1.8,y1=i[i$ringnumber=="2KZ09034"&i$reversals==2,]$learned,lty=2)

segments(x0=1.8,y0=i[i$ringnumber=="2KZ09034"&i$reversals==2,]$learned,

x1=2.8,y1=i[i$ringnumber=="2KZ09034"&i$reversals==3,]$learned,lty=2)

segments(x0=2.8,y0=i[i$ringnumber=="2KZ09034"&i$reversals==3,]$learned,

x1=3.8,y1=i[i$ringnumber=="2KZ09034"&i$reversals==4,]$learned,lty=2)

segments(x0=3.8,y0=i[i$ringnumber=="2KZ09034"&i$reversals==4,]$learned,

x1=4.8,y1=i[i$ringnumber=="2KZ09034"&i$reversals==5,]$learned,lty=2)

with(subset(i,i$ringnumber=="2KZ09034"),

points(x=reversals-0.2,y=learned,pch=c(4,1,4,1,4,1),col=as.character(color),cex=1.4))

**#Labels**

mtext("Batch 1 (October 2022)", outer=TRUE, cex=1.2,at = c(0.15))

mtext("Batch 2 (November 2022)", outer=TRUE, cex=1.2,at = c(0.40))

mtext("Batch 3 (January 2023)", outer=TRUE, cex=1.2,at = c(0.65))

mtext("Batch 4 (February 2023)", outer=TRUE, cex=1.2,at = c(0.90))
