## Supplementary figures and images for "Great tits show serial reversal learning in the perseverance phase but not in the new learning phase"

### Supplementary figure S1

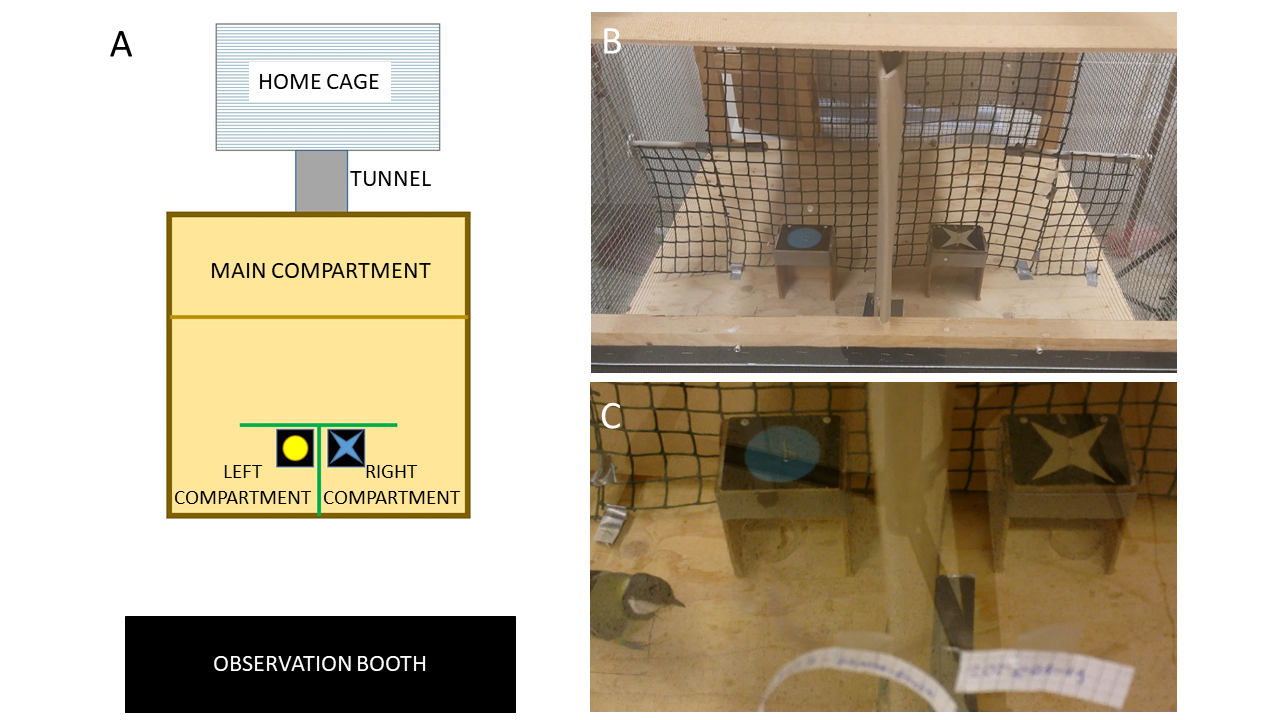

### Supplementary figure S2

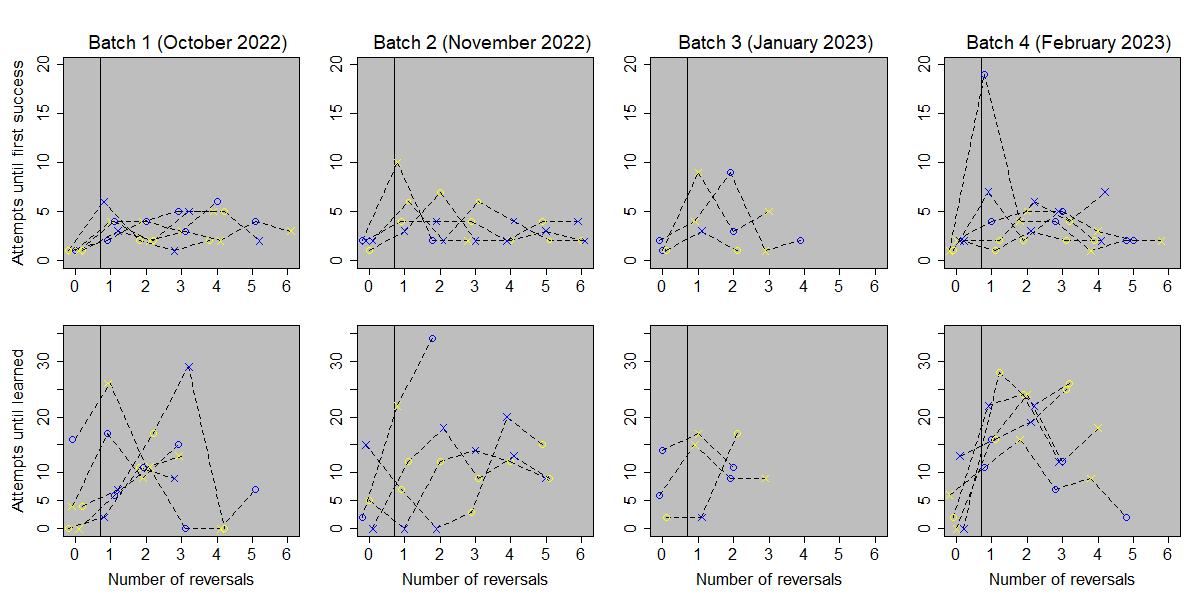
